# Distinct Motor Cortex Interneuron Plasticity and Its Association with Prefrontal Brain Volume in Parkinson’s Disease

**DOI:** 10.1101/2025.08.01.667976

**Authors:** Amanda O’Farrell, Nesrine Harroum, Amina El-Housseini, Pierre Blanchet, Alexandru Hanganu, Jason L. Neva

## Abstract

Parkinson’s disease (PD) is characterized by motor and cognitive deficits, including abnormal primary motor cortex (M1) excitability and diminished sensorimotor neuroplasticity. While paired associative stimulation (PAS) can induce M1 plasticity, people with PD (PwPD) demonstrate variability that cannot be accounted for by disease progression or medication status. Distinct M1 interneuron populations and attention-related brain structures may influence the reduced PAS- induced neuroplasticity. We aimed to characterize M1 interneuron plasticity in PwPD using attention-modulated PAS and identify neurostructural correlates. PwPD underwent MRI, then a PAS protocol with task-relevant attention. Transcranial magnetic stimulation (TMS) assessments of corticospinal excitability using posterior-to-anterior (PA) and anterior-to-posterior (AP) current directions were employed before and three post-PAS time-points. PAS induced distinct time- dependent M1 interneuron excitability changes. PA TMS showed increased corticospinal excitability at all post-PAS time-points; AP TMS increased only at 30 minutes. Rostral middle frontal gyrus volume uniquely explained variance in PA-sensitive M1 interneuron plasticity. In contrast, AP-sensitive plasticity was associated with baseline AP TMS excitability and age. These findings highlight that M1 interneuron circuits show unique neuroplasticity patterns in PwPD and relate to prefrontal brain volume. Our results suggest a complex interplay between motor and cognition-related deficits as interrelated pathophysiological features in PD.

## 1. Introduction

Parkinson’s disease (PD) is a progressive neurodegenerative disorder marked by motor (e.g., tremor, bradykinesia/akinesia) and non-motor (e.g., executive control dysfunction, impaired attention) signs and symptoms. PD-related signs and symptoms are linked to degeneration of the basal ganglia (i.e., substantia nigra), with other work demonstrating the critical role of basal ganglia-thalamocortical circuits in the known motor and cognitive functional impairments (Kalia & Lang, 2015; Popescu et al., 2024). Recent studies using transcranial magnetic stimulation (TMS) have also identified deficits in the excitability and neuroplasticity of sensorimotor cortices as important pathophysiological features of PD (Brown et al., 2014; Cantello et al., 1991; Ellaway et al., 1995; Wu et al., 2012). Specifically, people with PD (PwPD) exhibit inconsistent sensorimotor neuroplasticity in response to paired associative stimulation (PAS) (Bagnato et al., 2006; Kačar et al., 2013; Koch, 2013; Morgante et al., 2006; Ueki et al., 2006), which can induce transsynaptic long-term potentiation-like plasticity in the primary motor cortex (M1) (Carson & Kennedy, 2013; Stefan et al., 2000, 2004; Suppa et al., 2017).

Although there is evidence for diminished PAS-induced neuroplasticity in PwPD, the limited research has yielded conflicting results (Koch, 2013). Indeed, PwPD show reduced response to PAS compared to healthy controls, who typically demonstrate increased corticospinal excitability following the intervention (Morgante et al., 2006; Stefan et al., 2004; Ueki et al., 2006). However, other evidence demonstrates increased corticospinal excitability following PAS when PwPD are on dopaminergic medication and do not display dyskinesias (Morgante et al., 2006). Conversely, other work has shown that PwPD exhibited a greater increase in corticospinal excitability following PAS when “OFF” medication. In contrast, their increased response “ON” medication was akin to that of healthy controls (Bagnato et al., 2006). Still, other research showed impaired response to PAS in the more affected hemisphere and an increased response in the less affected hemisphere, specifically with newly diagnosed PwPD not currently on dopaminergic medication (Kojovic et al., 2012). These results suggest that other factors may play a critical role in the diminished neuroplasticity following PAS, which may be related to functional and structural differences within motor and non-motor brain regions that occur during PD progression (Hanganu et al., 2013; Kojovic et al., 2012, 2015; Sakato et al., 2024; Sokołowski et al., 2024). Supporting this notion, in advanced stages, research demonstrated diminished PAS response in PwPD who received dopaminergic medication for several years and those who did not receive medication, when compared to healthy controls (Kačar et al., 2013). These results suggest that dopaminergic medication does not consistently restore PAS-induced neuroplasticity, highlighting that there are likely other critical underlying mechanisms contributing to the variability of response to PAS (Kačar et al., 2013; Koch, 2013).

One potential contributor to the variability in sensorimotor neuroplasticity in PwPD is the critical role of attention allocation during PAS (Bagnato et al., 2006; Kačar et al., 2013; Morgante et al., 2006; Stefan et al., 2004). Critically, PAS-induced neuroplasticity has been shown to depend on directing task-relevant attention during the protocol, specifically by allocating attention to the limb receiving peripheral stimulation (Stefan et al., 2004). Importantly, PwPD frequently exhibit executive dysfunction and attentional deficits (Dirnberger & Jahanshahi, 2013; Kudlicka et al., 2011), specifically demonstrating diminished performance on tasks that require response inhibition (e.g., the Stroop test; (Bucur & Papagno, 2023; Dirnberger & Jahanshahi, 2013; Hamdan & Vieira, 2022) as well as sustained and selective attention (Cammisuli et al., 2021; Siepel et al., 2014). These deficits are also reflected in motor learning studies, where PwPD exhibit specific deficits in motor tasks that rely on attentional resources and cognitive strategies (Amick et al., 2006; Bédard & Sanes, 2011; Contreras-Vidal & Buch, 2003; Isaias et al., 2011; Marinelli et al., 2009, 2017; Moisello et al., 2015; Smiley-Oyen et al., 2006; Yamadori et al., 1996). Interestingly, deficits in motor learning were observed irrespective of medication status or disease severity (Bédard & Sanes, 2011; Isaias et al., 2011; Marinelli et al., 2009; Moisello et al., 2015). Taken together, these findings suggest that PwPD display diminished motor-related function compared to controls, particularly in tasks that require higher levels of cognitive strategy and attentional resources, which is consistent with research demonstrating frontal-striatal deficits (Auning et al., 2014; Hanganu et al., 2013; Lang et al., 2020; Monchi et al., 2004; Nagano-Saito et al., 2016). Despite the known importance of attention allocation for the induction of neuroplasticity via repetitive TMS protocols (Kamke et al., 2012; Stefan et al., 2004) and its impairment in PD (Auning et al., 2014; Hanganu et al., 2013; Lang et al., 2020; Monchi et al., 2004; Nagano-Saito et al., 2016), few studies have controlled attention allocation during PAS in PwPD (Kačar et al., 2017; Kojovic et al., 2012; Ueki et al., 2006). This leaves a significant gap in our understanding of how attentional factors, and thus prefrontal cortical functions, contribute to M1 neuroplasticity in PD.

A second potential contributor to the variability in PAS-induced neuroplasticity in PD relates to the neurophysiology and behaviour of distinct M1 interneuron populations that can be measured with different TMS current directions (Di Lazzaro et al., 2012; Hamada et al., 2014; Neva et al., 2021). While most TMS measurements during and following PAS typically employ a posterior-to-anterior (PA) current (Classen et al., 2004; Ni et al., 2019; Stefan et al., 2000, 2004), functionally distinct interneuron pools within M1 can be preferentially activated using an anterior-to-posterior (AP) TMS current (Di Lazzaro et al., 2001; Opie & Semmler, 2021). Single-pulse TMS generates multiple descending volleys of activity via direct (D-waves) or indirect (I-waves; early and late) activation of corticospinal output neurons in M1, resulting in observable motor evoked potentials (MEPs; (Di Lazzaro et al., 2001, 2012; Hamada et al., 2014; Hanajima et al., 1998, 2002; Ilić et al., 2002; Ziemann & Rothwell, 2000). At low intensities, PA and AP TMS preferentially activate early and late I-waves, respectively (Di Lazzaro et al., 1998, 2012). The patterns of I-waves elicited by these different TMS current directions likely represent distinct PA-sensitive and AP-sensitive interneuron circuits within M1 (Hamada et al., 2014; Hannah et al., 2018; Mirdamadi et al., 2017; Neva et al., 2021; Ni et al., 2019; Spampinato et al., 2020). Importantly, the excitability of these interneuron circuits is differentially modulated by motor tasks that require greater attention allocation and action selection (Hannah et al., 2018; Mirdamadi et al., 2017), as well as by adjunct interventions such as PAS (Hamada et al., 2014; Ni et al., 2019) and acute exercise (Neva et al., 2021). These results suggest that AP and PA-sensitive interneurons display distinct neuroplastic patterns and serve unique functional roles, acting as a potentially critical interface between prefrontal and motor-related brain regions (Hamada et al., 2014; Hannah et al., 2018; Mirdamadi et al., 2017). Thus, these interneurons may play a role in the diminished M1 neuroplasticity in PwPD. Critically, the role of AP- and PA-sensitive interneurons in PAS-induced M1 neuroplasticity remains unexplored in PwPD.

A third potential contributor to the variability in PAS-induced neuroplasticity may be the morphological changes in motor and non-motor-related regions characteristic of PD (Hanganu et al., 2014; Pieperhoff et al., 2022; Sokołowski et al., 2024). As PD is a neurodegenerative disease that typically develops later in life, individuals often face aging-related cortical thinning (Blinkouskaya et al., 2021). Additionally, brain volume atrophy is accelerated in PwPD, particularly in prefrontal cortical regions, precentral gyri and subcortical regions like the substantia nigra (Hanganu et al., 2014; Pieperhoff et al., 2022; Sokołowski et al., 2024). Given these structural differences and the known attention-related deficits in PwPD (Cammisuli et al., 2021; Gerrits et al., 2016; Hanganu et al., 2014; Wilson et al., 2019), prefrontal and motor-related brain region volume may contribute to M1 neuroplasticity following PAS (Chafee & Heilbronner, 2022; Eickhoff et al., 2016; Vo et al., 2023). However, the potential link between PAS-induced neuroplasticity and the volume of relevant prefrontal and motor-related brain regions remains unknown in PwPD.

Thus, in this study, we examined potential factors influencing the variability of neuroplasticity induced by PAS in PwPD. First, we aimed to characterize the modulation of M1 interneuron excitability following PAS in PwPD. We hypothesized that AP-sensitive interneuron excitability would be enhanced to a greater extent than PA-sensitive interneuron excitability following PAS, due to greater attention-related modulation of AP-interneuron excitability (Mirdamadi et al., 2017). Second, we aimed to identify the neurostructural correlates that may account for unique variance in PAS-induced M1 interneuron excitability change. We hypothesized that AP-sensitive interneuron excitability changes would be associated with prefrontal cortical volume, and to a lesser extent, M1 volume. We expected that PA-sensitive interneuron excitability change following PAS would be related to M1 cortical volume, and to a lesser extent, the prefrontal cortical regions.

## 2. Methods

### 2.1 Participants

Twenty-two individuals with mild to moderate PD were recruited (14 males; age range: 55–85 years; mean age: 69 ± 8.9). All participants were right-handed and reported being predominantly impaired on the left side. Mean Unified Parkinson’s Disease Rating Scale (UPDRS) scores were 32 ± 9.7 and taken while “ON” medication, and mean MOCA scores were 27 ± 2.2. Exclusion criteria included any contraindications to TMS or MRI. Informed consent was obtained before participation, and MRI safety screening was conducted in accordance with the guidelines of the Unité Fonctionnelle de Neuroimagerie (UNF). Additionally, TMS safety screening was performed. Six participants were excluded due to insufficient TMS responses or excessive muscle activity. The final sample included 15 participants who completed both the TMS and MRI sessions, with one additional participant completing only the TMS portion due to experiencing discomfort during the MRI. All participants were encouraged to continue taking their medication as usual. All procedures were approved by the Ethics Committee of the Centre de Recherche de l’Institut Universitaire de Gériatrie de Montréal (CRIUGM).

### 2.2. Experimental design

All participants took part in two experimental sessions, held on different days (Fig. 1). On the first day, MRI scans were taken, and participants completed the Edinburgh Handedness Inventory (Oldfield, 1971), the MoCA, and the UPDRS. On the second day, participants took part in the paired associative stimulation (PAS) protocol. Corticospinal excitability was assessed at four time points using both PA and AP TMS current directions: baseline (Pre), immediately post-(Post_0_), 15 minutes post- (Post_15_), and 30 minutes post-PAS (Post_30_). At each time point, motor evoked potentials (MEP) were recorded at 110% and 130% of resting motor threshold (RMT).

**Figure 1:**
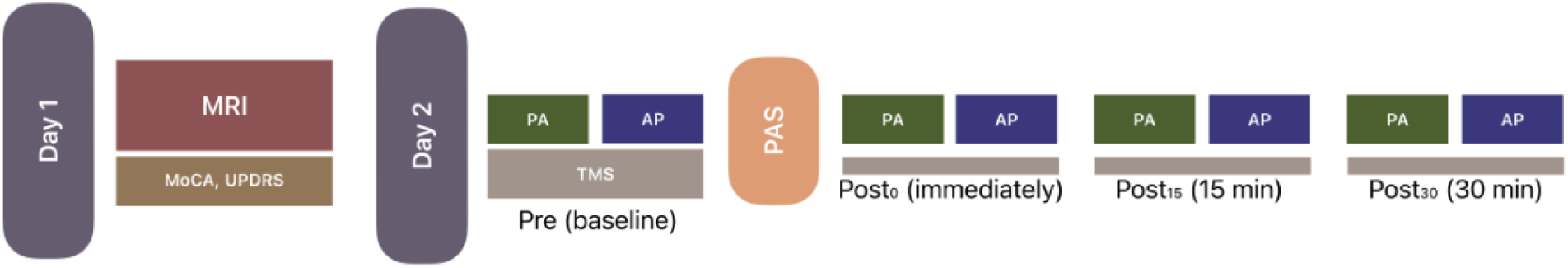
Study Design. A T1-weighted anatomical MRI scan was conducted on day 1 of testing. On the second day, measures of corticospinal excitability were taken using both anterior-posterior and posterior-anterior transcranial magnetic stimulation current directions. These measurements were taken before (Pre), immediately following (Post_0_), 15 minutes after (Post_15_), and 30 minutes after (Post_30_) an excitatory paired associative stimulation (PAS) protocol. <u>Abbreviations</u> AP: anterior-to-posterior; MoCA: Montreal Cognitive Assessment; MRI: magnetic resonance imaging; PA: posterior-to-anterior; PAS: paired associative stimulation; TMS: transcranial magnetic stimulation

### 2.3. MRI Acquisition and Preprocessing

MRI scans were acquired using a 3T Siemens Tim Trio MRI scanner with a 12-channel head coil. T1-weighted (T1w) anatomical scans were obtained with a field of view (FOV) of 256 × 256 mm, with a voxel size of 1 × 1 × 1 mm, a TR/TE of 2300/30 ms, and a flip angle of 9°. Cortical reconstruction and volumetric segmentation were performed using the Freesurfer image analysis package (Dale et al., 1999; Fischl et al., 1999). Precentral, postcentral, rostral middle frontal, and caudal middle frontal cortical volume values were extracted. The T1w images were also converted from DICOM to NIfTI format using MRIcroGL and stored in BIDS format. These files of the T1w scans were subsequently used in BrainSight (Rogue Research Inc., Montreal, QC, Canada) for neuronavigation during TMS assessments and PAS.

### 2.4. Electromyographic recording

MEPs were recorded from surface EMG electrodes placed over the right abductor pollicis brevis muscle (APB) and first dorsal interosseous (FDI) muscles using LabChart software (LabChart 8.0) and sampled using a PowerLab 26T (PL26T04 PowerLab, ADInstruments, Colorado Springs, CO, USA). Electrodes were arranged in a belly-tendon configuration over the APB and FDI, while the ground electrode was placed on the ulnar styloid. The APB was a muscle of primary interest, with the FDI included to record complementary data. Data were captured in a 500-ms sweep, from 100 ms before to 400 ms after TMS delivery, with an acquisition rate of 2 kHz, along with bandpass filtering (20-400 Hz) and notch filtering at a center frequency of 50 Hz.

### 2.5. Transcranial Magnetic Stimulation

Participants sat comfortably in an adjustable chair and remained at rest for all TMS measurements. A Magstim BiStim 200^2^ stimulator (Magstim Co., UK) connected to a figure-of-eight coil (Magstim 70 mm P/N 9790, Magstim Co., UK) delivered monophasic TMS pulses. All measurements before and after PAS were obtained using PA and AP TMS currents (Fig. 2) A standard coil was used for PA TMS to direct the current from the posterior to the anterior direction. A second custom coil was used for AP current direction TMS that delivered magnetic pulses in the opposite direction (180 degrees) compared to the PA coil. The TMS coils were positioned 45° to the mid-sagittal plane with the handle facing backward. Brainsight neuronavigation software (Rogue Research Inc., Montreal, QC, Canada) was used to ensure accurate and consistent coil position and monitoring. The M1 “hotspot” corresponding to the right abductor pollicis brevis (APB) was identified by placing the PA TMS coil over M1 using neuronavigation of individual MRI scans. The resting motor threshold (RMT) was determined at the hotspot for each TMS current direction (PA, AP). RMT was defined as the lowest stimulus intensity needed to elicit 5 out of 10 consecutive MEPs with a peak-to-peak amplitude of 50 μV or more. Throughout the study, TMS pulses were administered randomly between 0.15 and 0.2 Hz (with ∼20% variation).

**Figure 2:**
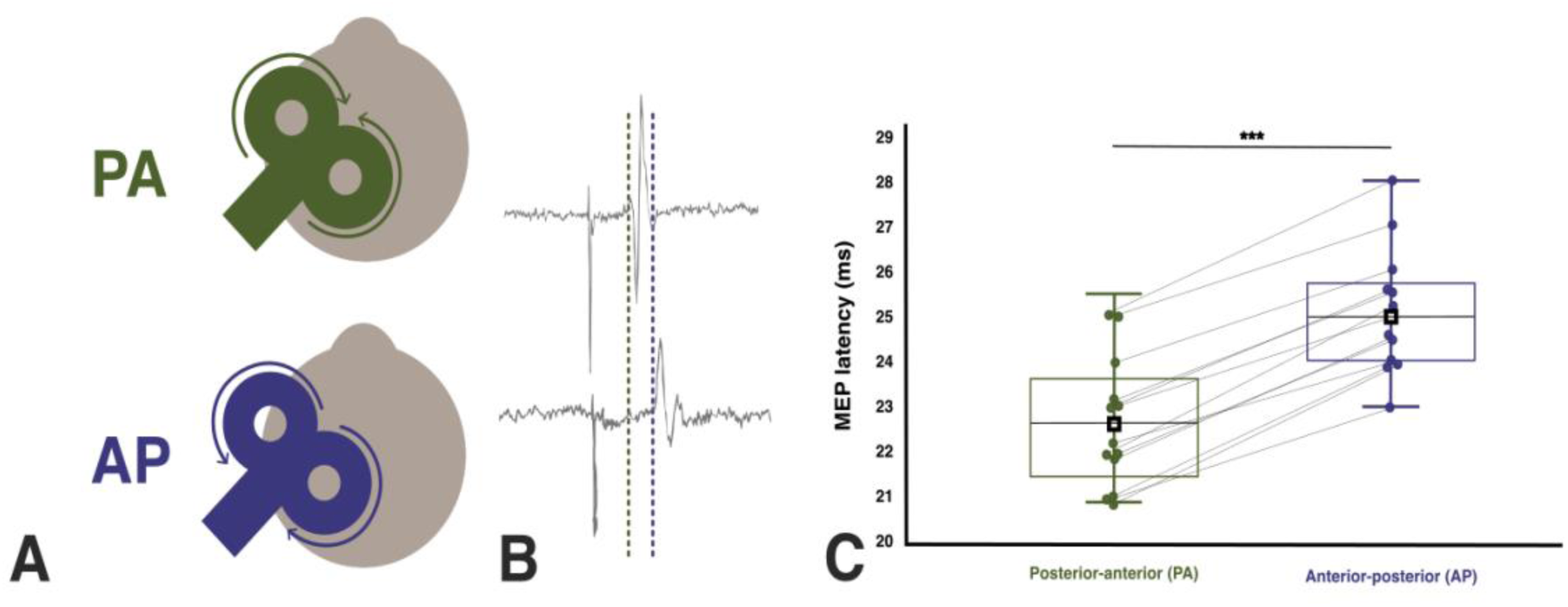
Transcranial magnetic stimulation (TMS) current directions and motor evoked potential (MEP) onset latency results. (A) TMS current directions are illustrated. PA TMS is shown above, and AP TMS is shown below. The figure illustrates the TMS coil current directions, represented by arrows, and orientations over the left M1 (dominant) and APB representations. We utilized a standard 8-figure TMS coil for PA stimulations and a reversed current coil for AP stimulations. The coil was oriented at 45° for both PA and AP TMS currents relative to the longitudinal fissure. (B) Electromyographic (EMG) traces by a representative participant were recorded from the right (dominant) APB. Vertical dashed lines indicate MEP onset latency elicited by each TMS current direction (PA, AP). (C) MEP onset latency for each TMS current (PA, AP) is shown using boxplots, with individual data points represented by dots and connected by grey lines. Black squares and lines across the boxes represent means. <u>Abbreviations</u>: AP: anterior-to-posterior; ms: milliseconds; PA: posterior-to-anterior; TMS: transcranial magnetic stimulation; *** p < .001.

#### Corticospinal excitability

MEP amplitudes were evaluated in both the PA and AP current directions to examine potential changes in corticospinal excitability following PAS. MEPs were assessed at 110% and 130% RMT in both PA and AP current directions. MEPs were assessed at 110% RMT, as previous work has shown that lower-intensity suprathreshold TMS pulses preferentially activate distinct interneuron circuits targeted with PA and AP currents in young, healthy participants (Cirillo et al., 2018; Di Lazzaro et al., 2001; Neva et al., 2021; Ni et al., 2019; Sale et al., 2016). Changes in MEP amplitudes were also assessed using 130% RMT, as TMS intensities in this range are conventionally used to assess changes in corticospinal excitability post-PAS (Bagnato et al., 2006; Morgante et al., 2006; Stefan et al., 2004). Fifteen stimuli were administered at each TMS intensity using both current directions (AP, PA), at all four different time points. Thus, the assessment of corticospinal excitability was performed by averaging the peak-to-peak MEP amplitudes of 15 trials at four time points: Pre (baseline), Post_0_ (immediately following PAS), Post_15_ (15 minutes following PAS), and Post_30_ (30 minutes following PAS).

#### MEP onset latencies

MEP onset latencies were identified as the earliest latency within a block of 15 MEPs. This method indirectly assessed I-wave recruitment and suggested preferential activation of specific interneuron populations based on the TMS current direction. The procedure was carried out for both PA and AP TMS current directions (measured at 110% RMT; Fig. 2) (Hamada et al., 2014; Neva et al., 2021; Ni et al., 2019; Sale et al., 2016; Spampinato, 2020).

### 2.6. Paired-Associative Stimulation (PAS)

The PAS protocol involved 180 stimulus pairs administered at a frequency of 0.25 Hz, with peripheral nerve stimulation preceding the TMS pulse by 25 ms (∼12 minutes total; Fig. 3) (Stefan et al., 2004). Stimulus pairs consisted of electrical peripheral stimulation of the right median nerve at 150% of the motor threshold (the minimum intensity that causes a visible APB twitch), followed by a PA TMS pulse applied over M1 at 130% RMT. The median nerve stimulation was administered using a constant-current electrical stimulator (DS7R; Digitimer, Welwyn Garden City, UK). To direct attention, an additional electrode was placed on the index finger of the right hand, delivering infrequent low-intensity electrical pulses via the PowerLab 26T isolated stimulator. During the PAS protocol, participants were instructed to attend to and count these stimuli, which occurred every ∼80 seconds and never simultaneously with the median nerve stimulation, with a total of 9 stimuli delivered during the PAS protocol. Participants were required to report the number of stimuli detected at the end of the PAS protocol (Stefan et al., 2004).

**Figure 3:**
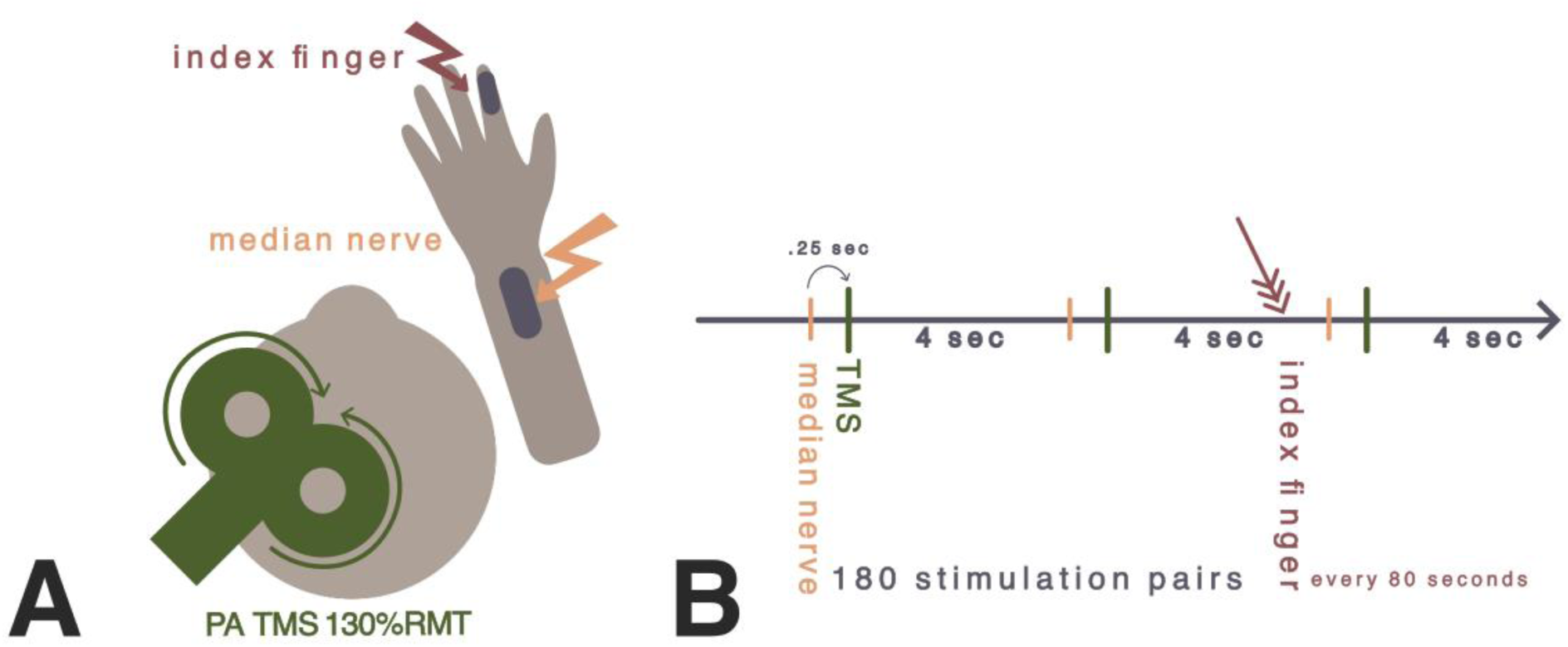
Paired-associative stimulation. (A) Protocol setup, PA TMS at 130% RMT over left M1. Two electrodes on the right upper limb, one over the median nerve and one over the right index finger. (B) PAS protocol timing with 25 ms between the peripheral nerve stimulation and PA TMS for 180 stimulation pairs. Attention was directed to the right hand by index finger stimulation occurring every ∼80 seconds. <u>Abbreviations</u>: sec: second; PA: posterior-to-anterior; RMT: resting motor threshold; TMS: transcranial magnetic stimulation.

### 2.7. TMS and MRI Data Processing

*TMS data*. Visual inspection was performed for all EMG data to detect voluntary muscle activity for all MEP trials. Peak-to-peak MEP amplitudes (measured in millivolts, mV) were analyzed using custom MATLAB scripts. Any trials showing visible voluntary pre-stimulus EMG activity were excluded from the analysis (5% of all trials).

#### MRI data

The Freesurfer processing pipeline involved T1 motion correction, removing non-brain tissue, performing automated Talairach transformation, segmenting subcortical white matter and deep gray matter structures, normalizing intensities, tessellating the gray-white matter boundary, applying automated topology correction, and deforming surfaces. Each brain was inflated to visualize the entire cortical surface. Gray matter sulco-gyral cortical units were automatically parcellated based on curvature landmarks (Destrieux et al., 2010) and labelled accordingly using conventional brain atlas nomenclature (Duvernoy, 2012). Regions of interest were restricted to the left hemisphere, which is the dominant and less affected hemisphere for our participants. They were intended to reflect hypothesized cortical regions associated with M1 plasticity induced by PAS. The sensorimotor cortices, encompassing the pre- and postcentral gyri, were included since M1 and the primary somatosensory cortex (S1) are directly involved in the PAS protocol, via TMS over M1 and median nerve stimulation relaying somatosensory information to S1 (Suppa et al., 2017). Both the rostral and caudal middle frontal areas were included because of their roles in cognitive functions, particularly attention allocation (Fig. 4).

**Figure 4:**
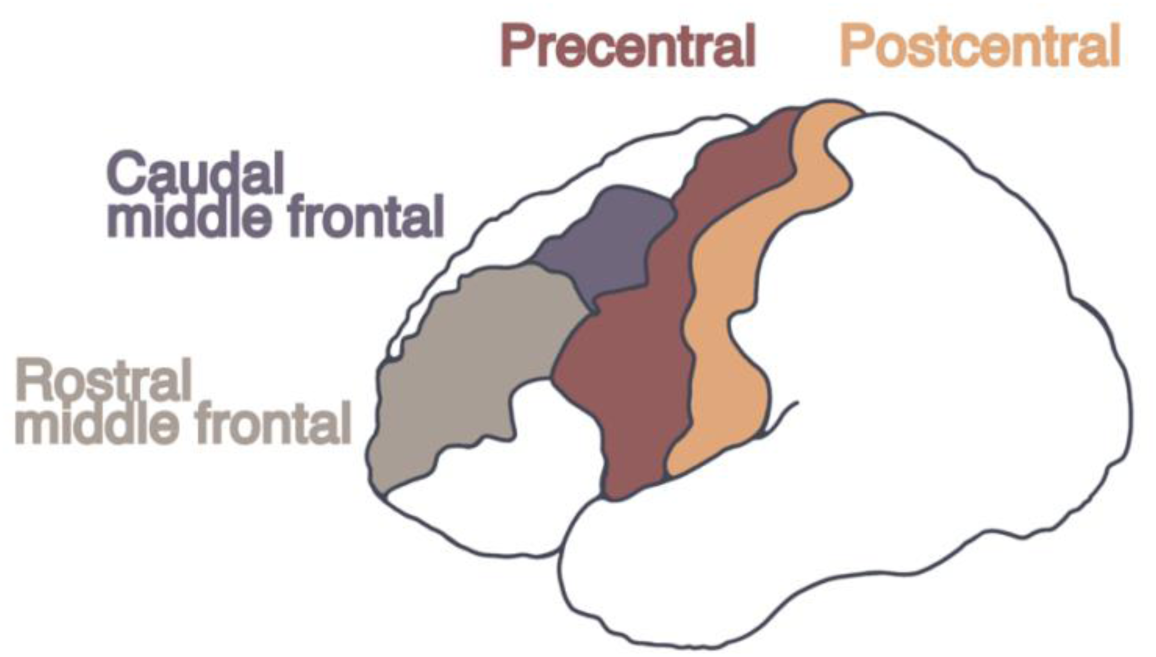
Regions of interest for brain volume analysis. Left hemisphere rostral middle frontal and caudal middle frontal are prefrontal cortical areas associated with cognition and attention. The precentral gyrus, also known as the primary motor cortex, is the area directly stimulated by TMS through PAS to induce neuroplasticity. It is also the cortical area where corticospinal excitability is measured using single-pulse TMS. The postcentral gyrus, the somatosensory area, receives peripheral sensory information in the cortex, and receives afferent signals dispatched from the peripheral nerve stimulation during PAS.

### 2.8. Statistical Analysis

To examine the effects of PAS on corticospinal excitability, analyses focused on changes in MEP amplitude from before (Pre) to the three post-PAS time points (Post_0_, Post_15_, Post_30_), as measured by the distinct TMS currents (PA, AP) and intensities (110%, 130% RMT). Two linear mixed models (one for each TMS current intensity: 110% and 130%RMT) were constructed to evaluate the main effects and interaction of TIME (Pre, Post_0_, Post_15_, Post_30_) and CURRENT (PA, AP), using participant as a random intercept and allowing for random slopes for TMS current directions. The models were estimated using the lme4 package in R (version 4.3.1), and Type III ANOVA with Satterthwaite’s method was used for significance testing (lmerTest). *Post hoc* analyses were conducted using the emmeans package with the Tukey correction where appropriate.

To assess whether the volumes of brain regions of interest accounted for variance in PAS-induced M1 interneuron excitability changes while considering age, MOCA scores, and baseline M1 excitability (i.e., RMT), two stepwise regression analyses were conducted using peak changes in corticospinal excitability post-PAS, measured with PA and AP TMS currents at 110% RMT. Specifically, peak changes post-PAS occurred at Post_15_, as measured with PA TMS, and at Post_30,_ as measured with AP TMS. Predictor variables included age, MOCA, RMT (RMT_PA_ or RMT_AP_, respectively), caudal middle frontal (CMF), rostral middle frontal (RMF), precentral and postcentral gyri. Left-hemisphere brain volumes were included, as participants were right-handed, and they corresponded to the employed PAS protocol.

Zero-order, part, and partial correlations were produced for each model. Acceptable collinearity between multiple predictors was determined using the variance inflation factor (below 10) and tolerance levels (above 0.1) (Field, 2005). Residual statistics and plots were generated to verify the normality and homoscedasticity of the data. Durbin-Watson diagnostics were used to ensure the independence of the residuals (Field, 2005). The significance level was set at p < .05 for each statistical test.

## 3. Results

### 3.1. PAS-induced M1 interneuron excitability change

#### MEP data at 110% RMT

A significant TIME by CURRENT interaction was found (*F_3, 1629.29_*= 4.705, *p* = .003; Fig. 5). For PA TMS current, *post hoc* analyses showed that corticospinal excitability increased at Post_0_ (*t_1635_* = -2.581, *p* = .049), Post_15_ (*t_1635_* = -5.001, *p* < .001), and Post_30_ (*t_1635_* = - 2.730, *p* = .032) compared to Pre. For AP TMS current, corticospinal excitability increased at Post_30_ relative to Pre (*t_1635_*= - 4.127, *p* < .001). Additionally, a significant main effect of TIME (*F_3, 1629.30_* = 9.9302, *p* < .001), but no main effect of CURRENT (*F_1, 15.47_* = 0.3048, *p* = .589), was found.

**Figure 5:**
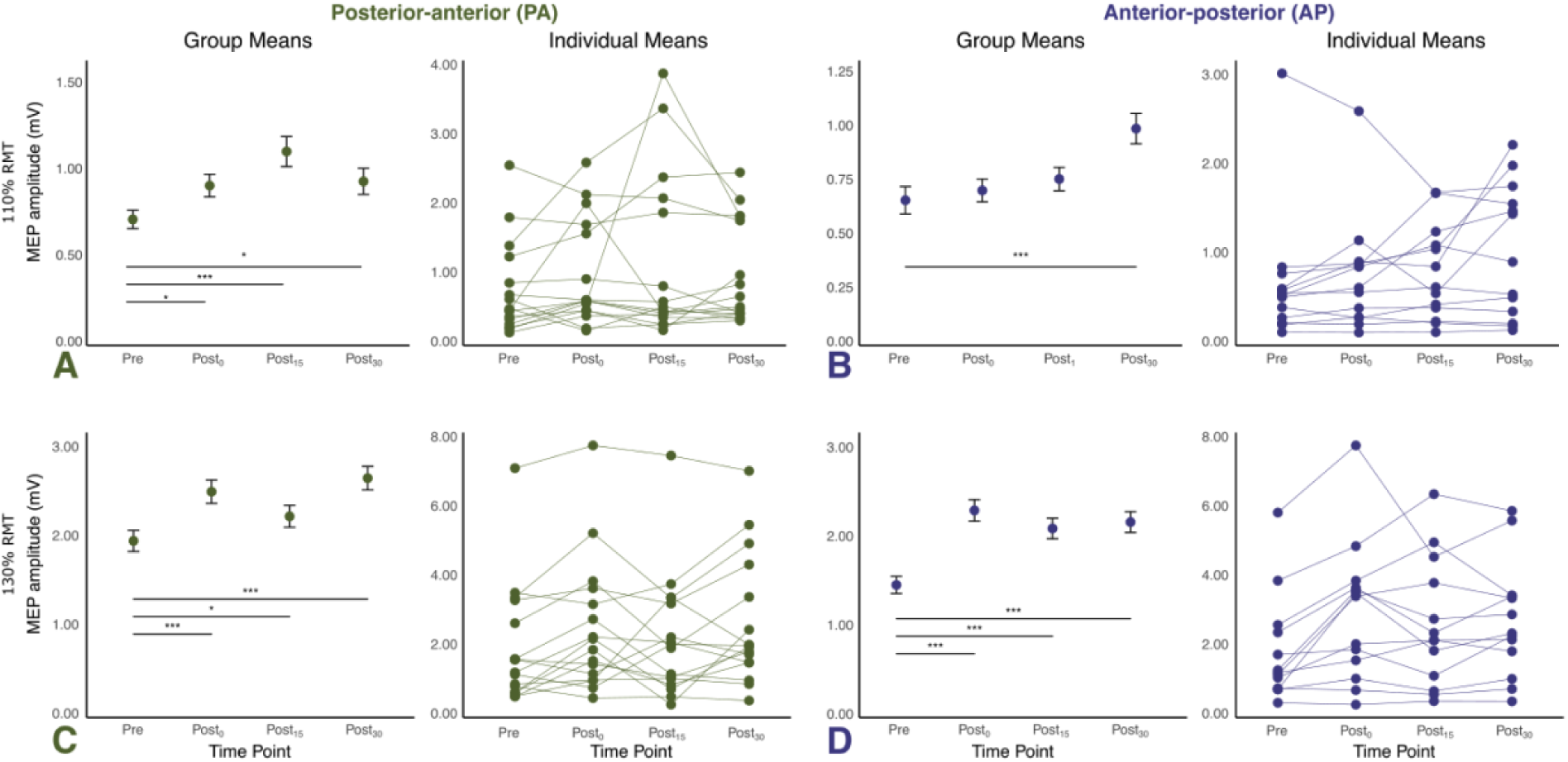
Change in MEP amplitude at 110% RMT and 130% RMT. (A, B) Display mean MEP amplitude at each time point with significant changes from baseline at 110% RMT and individual mean trajectories in the PA TMS current direction (A) and AP current direction (B). (C, D) Display mean MEP amplitude at each time point with significant changes from baseline at 130% RMT and individual mean trajectories in the PA TMS current direction (C) and AP current direction (D). <u>Abbreviations</u>: AP: anterior-to-posterior; MEP: motor evoked potential; mV: millivolt; PA: posterior-to-anterior; RMT: resting motor threshold.

#### MEP data at 130% RMT

A significant two-way interaction was found between TIME and CURRENT (*F*_3, 1598.26_ = 3.035, *p* = .028). For PA TMS current, *post hoc* analyses revealed an increase in corticospinal excitability at Post_0_ (*t_1604_* = -5.439, *p* < .0001), Post_15_ (*t_1604_* = -2.632, *p* = .043), and Post_30_ (*t_1604_* = -6.570, *p* = < .0001) compared to Pre. For AP TMS current, corticospinal excitability significantly increased at Post_0_ (*t_1604_* = -7.587, *p* < .0001), Post_15_ (*t_1604_* = -5.553, *p* < .0001), and Post_30_ (*t_1604_* = -6.240, *p* = < .0001) compared to Pre. We also found a significant main effect of TIME (*F*_3, 1598.26_ = 37.448, *p* < .001), but not CURRENT (*F_1, 15.51_* = 0.927, *p* = .350).

### 3.2. Regression modelling of PAS-induced M1 interneuron neuroplasticity

Stepwise regression analysis revealed that RMF brain volume accounted for a significant proportion of the variance in peak PA interneuron excitability change post-PAS (ΔR² = 0.390, *p* = .013; results and plots are shown in Table 1 and Figure 6). The model that included only RMF brain volume explained the greatest amount of unique variance in M1 excitability change assessed with PA TMS and was significant (R^2^ = 0.343; F_1,14_ = 8.300; *p* = .013), with no other brain volume (CMF, precentral and postcentral), nor age, MOCA or RMT_PA_ contributing to the model.

**Figure 6:**
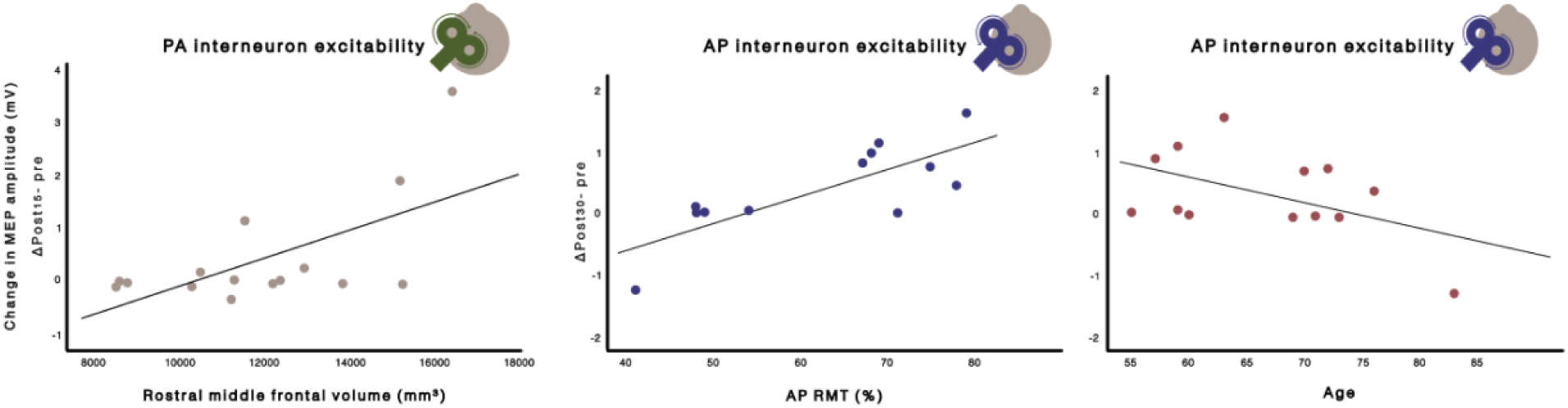
Bivariate correlations for peak significant change from Post_15_-Pre for PA TMS and Post_30_-Pre for AP TMS. <u>Abbreviations</u>: AP: anterior-to-posterior; MEP: motor evoked potential; mV: millivolt; PA: posterior-to-anterior; RMT: resting motor threshold.

**Table 1.** Linear regression models examining the associations between left-hemisphere cortical volumes and peak corticospinal excitability change post-PAS (PA and AP). RMF = rostral middle frontal.

| Variable | B | SE | $\beta$ | $t$ | $p$ | 95%CI |
| --- | --- | --- | --- | --- | --- | --- |
| ***PA TMS current corticospinal excitability change: $\Delta\text{Post}_{15}$-Pre*** |  |  |  |  |  |  |
| (Constant) | -2.73 | 1.11 |  | -2.45 | 0.03 | [-5.14, -0.32] |
| RMF volume | 0.00 | 0.00 | 0.62 | 2.881 | 0.01 | [0.00, -0.00] |
| R<sup>2</sup> | .39 |  |  |  |  |  |
| Adjusted R<sup>2</sup> | .34 |  |  |  |  |  |
| F<sub>1, 13</sub> | 8.30 |  |  |  |  |  |
| p(overall) | 0.013 |  |  |  |  |  |
| N | 15 |  |  |  |  |  |
| ***AP TMS current corticospinal excitability change: $\Delta\text{Post}_{30}$-Pre*** |  |  |  |  |  |  |
| (Constant) | -0.03 | 1.084 |  | -0.03 | 0.98 | [-2.50, 2.44] |
| AP RMT | 0.04 | 0.01 | 0.76 | 4.67 | < 0.01 | [0.020, 0.060] |
| Age | -0.03 | 0.01 | -0.39 | -2.40 | 0.04 | [-0.062, -0.002] |
| R<sup>2</sup> | 0.76 |  |  |  |  |  |
| Adjusted R<sup>2</sup> | 0.71 |  |  |  |  |  |
| F<sub>2, 9</sub> | 14.60 |  |  |  |  |  |
| P(overall) | .001 |  |  |  |  |  |
| N | 12 |  |  |  |  |  |

A second stepwise regression analysis revealed that RMT_AP_ (ΔR^2^ = 0.614, *p* = .003) and age (ΔR^2^ = 0.151, *p* = .040) accounted for significant and unique variance in peak AP interneuron excitability change post-PAS, resulting in a model that explained the greatest amount of variance in this change (R^2^ = 0.712; F_1,11_ = 14.598; *p* = .001; results and plots shown in Table 1 and Figure 6). Contrary to our hypothesis, none of the brain region volumes accounted for variance in AP M1 excitability change after PAS.

## 4. Discussion

This study investigated how PAS with task-relevant attention impacts M1 interneuron excitability in PwPD. Our main findings highlight that PwPD demonstrate distinct patterns of M1 interneuron plasticity. Specifically, we observed a time-dependent modulation of excitability in both PA- and AP-sensitive M1 interneuron populations following PAS. Additionally, PA-sensitive M1 excitability changes were found to be related to prefrontal brain volume, while AP-sensitive changes were associated with age and baseline M1 excitability. These findings suggest that the capacity for M1 plasticity in PwPD involves a complex interplay between motor and cognitive domains, and the deficits observed within them underscore potentially interrelated pathophysiological features of PD. Therefore, we will focus the following discussion on neurophysiological and behavioural mechanisms related to our findings.

### 4.1. Distinct patterns of M1 interneuron plasticity in people with PD

Our findings reveal distinct temporal patterns of attention-modulated PAS-induced plasticity in PA- and AP-sensitive interneurons in PwPD. Specifically, all post-PAS timepoints showed increased corticospinal excitability measured with PA TMS. Further, the peak increase occurred at 15 minutes post-PAS, which was earlier than that measured with AP TMS. Several previous studies have demonstrated inconsistent results following PAS in PwPD. Ueki et al. (2006) reported increased corticospinal excitability immediately following PAS, but only in participants who were “ON” dopaminergic medication. Similarly, Morgante et al. (2006) found that PAS-induced plasticity was restored when “ON” medication, nevertheless demonstrating inconsistent increases in corticospinal excitability post-PAS (Morgante et al., 2006; Ueki et al., 2006). Critically, PwPD displaying dyskinesias did not show increased corticospinal excitability post-PAS. In contrast, Kojovic et al. (2012) showed that newly diagnosed PwPD who had not yet taken dopaminergic medication displayed an increase in corticospinal excitability post-PAS on their less affected side, which was not found on their more affected side. We found a consistent increase in corticospinal excitability on the less affected side up to ∼40 min post-PAS, which aligns with some previous findings in PwPD taking dopaminergic medication (Morgante et al., 2006; Ueki et al., 2006). Critically, our results differ from other research showing either no increase in corticospinal excitability following PAS (Ueki et al., 2006) or a diminished response (Kačar et al., 2013; Morgante et al., 2006). This distinction may be due to the control of task-relevant attention to the target hand during PAS. Task-relevant attention directed toward the target hand has been shown to be critical for PAS-induced neuroplasticity in healthy individuals (Stefan et al., 2004). Although previous studies have instructed participants to “pay attention” to the target hand (e.g., by counting every other peripheral nerve stimulus applied during PAS (Kojovic et al., 2012)), this may have become redundant and difficult to attend to, particularly due to the high number of peripheral nerve stimulations and the lack of an engaging task during PAS. In our study, we closely followed the attention-direction protocol described in Stefan et al. (2004), which applied a separate weak electrical stimulation, which was infrequent (in our case, a total of 9 stimuli) and asynchronous to the TMS and median nerve stimulation (Stefan et al., 2004). Importantly, these particular parameters were previously chosen to enhance the ability to direct task-relevant attention to the unique stimuli being applied to the index finger, which totalled a number that did not overload working memory capacity (Oberauer et al., 2016; Stefan et al., 2004; Wilhelm et al., 2013), thereby allowing the participant to be more engaged in the attention task during PAS. Thus, the attention-related parameters employed during our PAS protocol may have contributed to our consistent and robust increase in corticospinal excitability post-PAS. Relatedly, our findings demonstrating an association with prefrontal brain volumes support the notion that the efficient engagement of attention-related brain circuits during our PAS protocol may have contributed to a more consistent post-PAS increase in corticospinal excitability.

The assessment of unique M1 interneurons using PA and AP TMS currents may have contributed to our findings of PAS-induced neuroplasticity in PwPD. There is a growing body of evidence that PA- and AP-directed TMS currents over M1 can preferentially activate distinct sets of interneuron inputs to corticospinal output neurons (Day et al., 1989; Di Lazzaro et al., 2001; Hamada et al., 2014; Hannah et al., 2018; Mirdamadi et al., 2017; Neva et al., 2021; Ni et al., 2019; Sale et al., 2016). Recent work has shown PAS differential changes in corticospinal excitability as measured with PA and AP TMS in healthy adults (Hamada et al., 2014; Ni et al., 2019). Specifically, Ni et al. (2019) found that PAS employed with a PA current increased corticospinal excitability measured with AP TMS current. In contrast, this effect was only a trend for an increase when measured with PA TMS current (Ni et al., 2019). Further, Hamada et al. (2014) found that subthreshold PAS with a 25 ms ISI requires AP current to elicit a robust corticospinal excitability increase, whereas PAS with a 21.5 ms ISI requires PA TMS (Hamada et al., 2014). These findings suggest that unique patterns of neuroplasticity of PA- and AP-sensitive interneuron populations can be induced with repetitive TMS protocols (e.g., PAS) in healthy adults (Hamada et al., 2014; Ni et al., 2019). However, the current study is the first to investigate this in PwPD. Here, we extend previous findings by demonstrating unique changes in corticospinal excitability post-PAS via assessment with PA and AP TMS currents. These results suggest that PA- and AP-sensitive M1 interneuron excitability highlight distinct pathophysiological features of PD, which are discussed below.

We found increased AP-sensitive corticospinal excitability uniquely at 30 minutes following PAS, occurring at a delay compared to PA-sensitive changes. These AP-sensitive interneurons represent a unique population within M1 that may play a crucial role in various functions highly relevant to PD, including attentional load management, motor preparation and adaptation, as well as neurophysiological connectivity with brain regions (e.g., the cerebellum) implicated in PD-related deficits in motor function (Hamada et al., 2014; Hannah et al., 2018; Mirdamadi et al., 2017). Previous work has shown that attentional load modulates AP-, but not PA-sensitive, interneuron excitability, suggesting a critical role in attention regulation during motor-related tasks (Mirdamadi et al., 2017). Since PwPD display deficits in sustained and selective attention and diminished performance at motor tasks requiring higher levels of attention and executive control (Cammisuli et al., 2021; Marinelli et al., 2009, 2017; Pilgrim et al., 2021), AP-sensitive interneurons may be associated with performance deficits in such motor tasks. However, this is beyond the scope of the current study, and future work should consider the association of corticospinal excitability assessed with PA and AP TMS currents with motor and cognitive deficits related to PD.

### 4.2. Prefrontal brain volume association with PA-sensitive M1 interneuron plasticity

A key finding of this study is that the volume of the rostral middle frontal gyrus (RMF) accounts for unique variance in PA-sensitive M1 interneuron plasticity. The volume of the RMF gyrus was positively related to increased PA-sensitive M1 interneuron plasticity, indicating that larger brain volume was associated with a greater increase in corticospinal excitability following PAS. The involvement of the rostral middle frontal cortex in this effect suggests the crucial role of cognitive processes, such as attention, in M1 interneuron excitability modulation. The RMF is widely recognized for its involvement in executive control, sustained attention, and working memory (Bonelli & and Cummings, 2007; Vincent et al., 2008). Relatedly, our findings align with previous work in healthy individuals, which revealed that task-relevant attention directed to the hand involved in PAS was critical to increased corticospinal excitability (Stefan et al., 2004). We extend these results to PwPD by showing that this PAS-induced neuroplasticity, involving task-relevant attention, may rely partly on preserved prefrontal cortical volume. Thus, our findings provide a partial explanation for inconsistent findings of PAS-induced neuroplasticity in PwPD (Bagnato et al., 2006; Kačar et al., 2017; Koch, 2013; Kojovic et al., 2012; Morgante et al., 2006; Ueki et al., 2006).

Contrary to our hypothesis, we did not observe an association of PAS-induced change in corticospinal excitability with precentral or postcentral brain volumes. Previous work has shown that the cortical volume of the region targeted by TMS is associated with the behavioural or neurophysiological response to stimulation in individuals with chronic stroke (Borich et al., 2015; Brodie et al., 2014). Specifically, it has been shown that enhanced motor learning in individuals with stroke following repetitive TMS to ipsilesional S1 was associated with greater brain volume of the post-central gyrus (Brodie et al., 2014). We have also demonstrated that greater M1 volume is associated with increased corticospinal excitability, as measured by TMS, in individuals with stroke (Borich et al., 2015). Thus, we expected an association of S1 and M1 brain volumes with PAS-induced neuroplasticity, as it involves the integration of somatosensory input and motor cortex output, which can be impaired in PwPD (Nelson et al., 2018; Stefan et al., 2000). Our results suggest that brain structures involved in attention allocation and cognition play a crucial role in sensorimotor neuroplasticity in PwPD, which aligns with the known motor and cognition-related deficits in this population (Chu et al., 2024; Cristini et al., 2023; Kalia & Lang, 2015; Siepel et al., 2014).

### 4.3. Age and RMT_AP_ association with AP-sensitive M1 interneuron plasticity

We found that AP-sensitive M1 interneuron plasticity relates to age and RMT_AP_ (i.e., baseline cortical excitability). Contrary to our hypothesis, we did not find an association of AP-sensitive plasticity with prefrontal and motor-related brain volumes. Specifically, we found that increasing age was related to less AP-sensitive neuroplasticity. This result aligns with findings on the impact of aging on the brain, where neuroplastic potential decreases with age (Fathi et al., 2010; Opie et al., 2017; Rogasch et al., 2009; Todd et al., 2010). Importantly, PD is an aging-related progressive neurodegenerative condition, which typically develops later in life (Collier et al., 2017; Levy, 2007). Other research indicates that AP-sensitive M1 excitability demonstrates unique aging-related characteristics (Sale et al., 2016). Specifically, short-interval intracortical inhibition (SICI), measured using AP TMS current, exhibited age-related differences, with older individuals displaying increased inhibition. This age-related difference was not apparent when using a PA TMS current (Sale et al., 2016). Thus, it is possible that with increasing age, SICI measured with AP TMS current may be enhanced, reducing responsiveness to a plasticity-inducing protocol like PAS (Kujirai et al., 2006; Ziemann et al., 2008). Regardless, unique aging-related changes in cortical excitability measured with AP and PA TMS currents may partly explain the findings, though several other potential contributing factors may be at play. Yet, this is speculative and requires further research to uncover the aging-related and PD-related impact on M1 interneuron plasticity.

Our results showed an association between changes in AP-sensitive interneuron excitability and RMT_AP_ (i.e., baseline M1 excitability). Specifically, a higher RMT_AP_ (% maximum stimulator output) value was associated with a greater increase in AP-sensitive interneuron excitability following PAS. Although this finding may be initially counterintuitive, it could suggest a "floor" effect, where individuals with lower excitability have more potential for excitability change following PAS. Yet, it is curious why this would only be the case for AP-sensitive interneuron excitability, requiring further research to address this question. Alternatively, since AP-sensitive interneurons typically have a higher threshold compared to PA-sensitive interneurons (Di Lazzaro et al., 2001; Hamada et al., 2014), they may have a greater potential for excitability change when in a less excitable state. This may be indicative of a compensatory neurophysiological characteristic in PwPD. Yet, this is speculative and requires further research to disentangle the potential explanations for our association.

### 4.4. Limitations

This study had some limitations. First, sample size should be taken into consideration when interpreting findings. Although we recruited a total of 22 PwPD to participate in the study, the final sample was reduced due to resting tremor and involuntary background EMG activity during TMS data collection, as well as a participant who experienced claustrophobia during the MRI scan. Despite the difficulties in initial recruitment and complications during data collection, the present results offer insights into the contributors to sensorimotor neuroplasticity as a pathophysiological feature of PD, which should be confirmed in future studies with larger sample sizes. Second, this study examined PwPD while “ON” dopaminergic medication. Thus, we did not include a second session when individuals were ‘OFF’ medication. This decision was based on practical considerations, including participant discomfort during laboratory visits, and to minimize challenges to participant recruitment and adherence. Finally, we did not include an additional attention condition during PAS (e.g., task-irrelevant attention allocation), which has been shown to suppress the expected enhancement in corticospinal excitability following PAS (Stefan et al., 2004). An ideal design would have included this PAS condition to test the potential suppressive effect of task-irrelevant attention on M1 plasticity in PD, thereby further confirming the critical role of attention-related circuits in M1 neuroplasticity in PwPD. Our reason for the decision not to include this condition was twofold: (i) to mitigate challenges with participant recruitment and adherence in the study, and (ii) to prioritize the investigation of the impact of task-relevant attention allocation during PAS on neuroplasticity of distinct interneuron pools in M1. Future studies should explore a broader range of attentional states further to understand the impact of PAS-induced neuroplasticity in PwPD.

## 5. Conclusion

This study found that PwPD exhibit distinct patterns of M1 interneuron plasticity following PAS while directing task-relevant attention. Specifically, we observed distinct changes in M1 interneuron excitability post-PAS as measured by PA and AP TMS currents. Peak PA interneuron excitability change was associated with prefrontal brain volume. In contrast, AP interneuron excitability change was associated with increased age and decreased baseline M1 excitability. These findings provide further evidence that PA and AP TMS can assess distinct M1 interneuron circuits, which show differential excitability changes after PAS in PwPD. Overall, these findings suggest that M1 plasticity in Pw PD involves a complex interaction between motor and cognitive domains, where deficits in these areas highlight potentially interconnected pathophysiological features of PD.

## Data Availability Statement

All main data presented in this manuscript are included within the figures. The datasets produced and analyzed in this study are available from the corresponding author upon reasonable request.

## Author Contributions

AOF contributed to the recruitment of PwPD, collected and analyzed the data and wrote the first draft of the manuscript. NH contributed to data collection. AEH contributed to participant recruitment and data collection. PB and AH contributed to participant recruitment and to editing the manuscript. AH contributed to the conception of the study and the experimental design. JLN conceived of the project, contributed to the interpretation of data, as well as writing and editing the manuscript. All authors edited the manuscript and approved the final version before submission.

## Acknowledgements

We would like to thank the students and research assistants who contributed to this work, and our participants for their time and commitment to taking part in this study.

